# Single-Session HRV Biofeedback Alters Neural Activity but Not Subjective Anxiety

**DOI:** 10.64898/2026.11.10.765768

**Authors:** Claire Dao, Isabella Man, Kate Healy, Kristen Estel, Stephanie Nelli

## Abstract

Anxiety and stress are among the most prevalent mental health concerns worldwide, yet barriers to effective treatment remain high. Biofeedback-based approaches offer a promising, low-cost complement to traditional interventions. The present study examined whether real-time heart rate variability (HRV) biofeedback enhances the efficacy of paced breathing for reducing state anxiety, and whether self-reported and neural measures of anxiety relief converge. Healthy adults completed a speeded Stroop task for stress induction, followed by a ten-minute paced breathing exercise. Participants were randomly assigned to an HRV biofeedback condition, in which real-time cardiac variability was displayed, or a control condition with the same breathing cue and no physiological feedback. State anxiety was assessed using the Spielberger State-Trait Anxiety Inventory (STAI-S), and neural activity was measured via EEG spectral power across delta, theta, alpha, and beta frequency bands. The Stroop task reliably elevated state anxiety, and paced breathing reduced STAI-S scores equivalently in both groups; HRV biofeedback did not confer additional subjective benefit. In contrast, EEG analyses revealed that the biofeedback group showed increases in delta and bilateral theta power relative to controls, and spatially widespread increases in alpha and beta power. Time-frequency analyses further showed more extensive and sustained spectral modulation in the biofeedback condition. These findings reveal a dissociation between self-reported and neural responses to a brief HRV biofeedback intervention: paced breathing alone was sufficient to reduce acute state anxiety, while HRV biofeedback produced distinct changes in neural oscillatory activity without enhancing self-reported anxiety relief. These results show that subjective anxiety recovery and neural recovery can diverge during acute HRV biofeedback, suggesting that physiological engagement with biofeedback emerges before conscious benefits.

## INTRODUCTION

Anxiety and stress disorders, among the most prevalent mental health concerns globally, are associated with alterations in subjective experience and physiological regulation, including changes in autonomic nervous system function, cardiovascular activity, and neural dynamics^1–6^. Although effective treatments exist, barriers including cost, accessibility, and stigma continue to motivate interest in low-cost, scalable interventions that can be delivered outside traditional clinical settings. Among these, paced breathing and heart rate variability (HRV) biofeedback have received considerable empirical attention as methods for improving autonomic regulation and reducing anxiety-related symptoms.

Biofeedback provides individuals with real-time information about physiological processes that typically operate outside conscious awareness, such as heart rate, respiration, and skin conductance, aiming to facilitate self-regulation of autonomic function^7,8^. HRV reflects variation in the timing between successive heartbeats and is widely regarded as an index of autonomic flexibility and parasympathetic regulation^9–13^. When combined with slow-paced breathing, a technique that increases HRV through enhanced cardiorespiratory coupling and vagal activity^14,15^, biofeedback has been shown to reduce self-reported anxiety across clinical and non-clinical populations^16–19^.

Much of the research on anxiety on self-report. Subjective ratings provide valuable information about conscious emotional experience^20,21^, but may not fully capture underlying physiological or neural processes^22^. Individuals often show imperfect awareness of their own internal physiological state, and self-reports may be influenced by expectation, response bias, or demand characteristics, to name a few^23,24^. Electroencephalography (EEG) offers complementary, millisecond-resolution measurement of oscillatory signatures linked to anxiety^25–27^. For example, heightened arousal and anxiety are frequently associated with beta activity^25,28^, whereas relaxation and internally directed processing are often characterized by increased alpha and theta oscillations respectively^29–31^. So, frequency-specific signatures may provide a useful framework for examining the neural consequences of biofeedback-based anxiety interventions and determining whether physiological and subjective indices converge.

Despite interest in HRV biofeedback, relatively few studies have simultaneously evaluated subjective anxiety and neural activity during acute biofeedback interventions. This question is important for understanding mechanisms, particularly during brief exposures where subjective and physiological responses may evolve on different timescales. The present study investigated the effects of a single-session HRV biofeedback intervention following experimentally induced stress. Healthy adults completed a speeded Stroop task to elevate state anxiety and were then randomly assigned to a ten-minute paced breathing exercise with or without real-time HRV biofeedback using the Polar H10 chest-strap monitor^32,33^. Anxiety was assessed using both the STAI-S and EEG spectral power. We predicted that paced breathing would reduce state anxiety in both groups and that HRV biofeedback would further enhance these effects. Our results reveal instead a meaningful dissociation: HRV biofeedback produced distinct changes in EEG oscillatory activity without providing additional reductions in self-reported anxiety. These findings suggest that neural and subjective indices of anxiety recovery may be partially dissociable during brief biofeedback interventions, and they highlight the value of incorporating objective neural measures when evaluating psychophysiological treatments.

## RESULTS

### State Anxiety (STAI-S)

State anxiety was assessed at three phases: baseline (upon arrival), stress (following the Stroop task) and recovery (following the breathing intervention) (Fig. 1). Mean STAI-S scores at baseline were 35.62 ± 9.60 (mean ± SD) across all participants and did not differ between groups (Fig. 2) (one-way ANOVA: *F*_1,35_ = 0.94, *p* = 0.34). STAI scores increased after the Stroop task (*t*_36_ = 3.97, *p* = 0.0003; 41.16 ± 9.46 mean ± SD), confirming successful stress induction. Scores then decreased during recovery (*t*_36_ = −4.44, *p* < 0.0001) to levels statistically equivalent to pre-task baselines (*t*_36_ = −1.65, *p* = 0.1; 33.83 ± 10.08, mean ± SD), indicating complete subjective recovery. A two-way repeated-measures ANOVA confirmed a significant main effect of phase (*F*_2,70_ = 3.52, *p* = 0.035) no significant main effect of biofeedback condition (*F*_1,35_ = 1.14, *p* = 0.29), and no significant condition-by-time interaction (*F*_2,70_ = 0.44, *p* = 0.64), reflecting mixed results in prior single-session biofeedback studies^34,35^. These results indicate that paced breathing reduced subjective state anxiety equivalently regardless of whether HRV biofeedback was provided.

**Figure 1.**
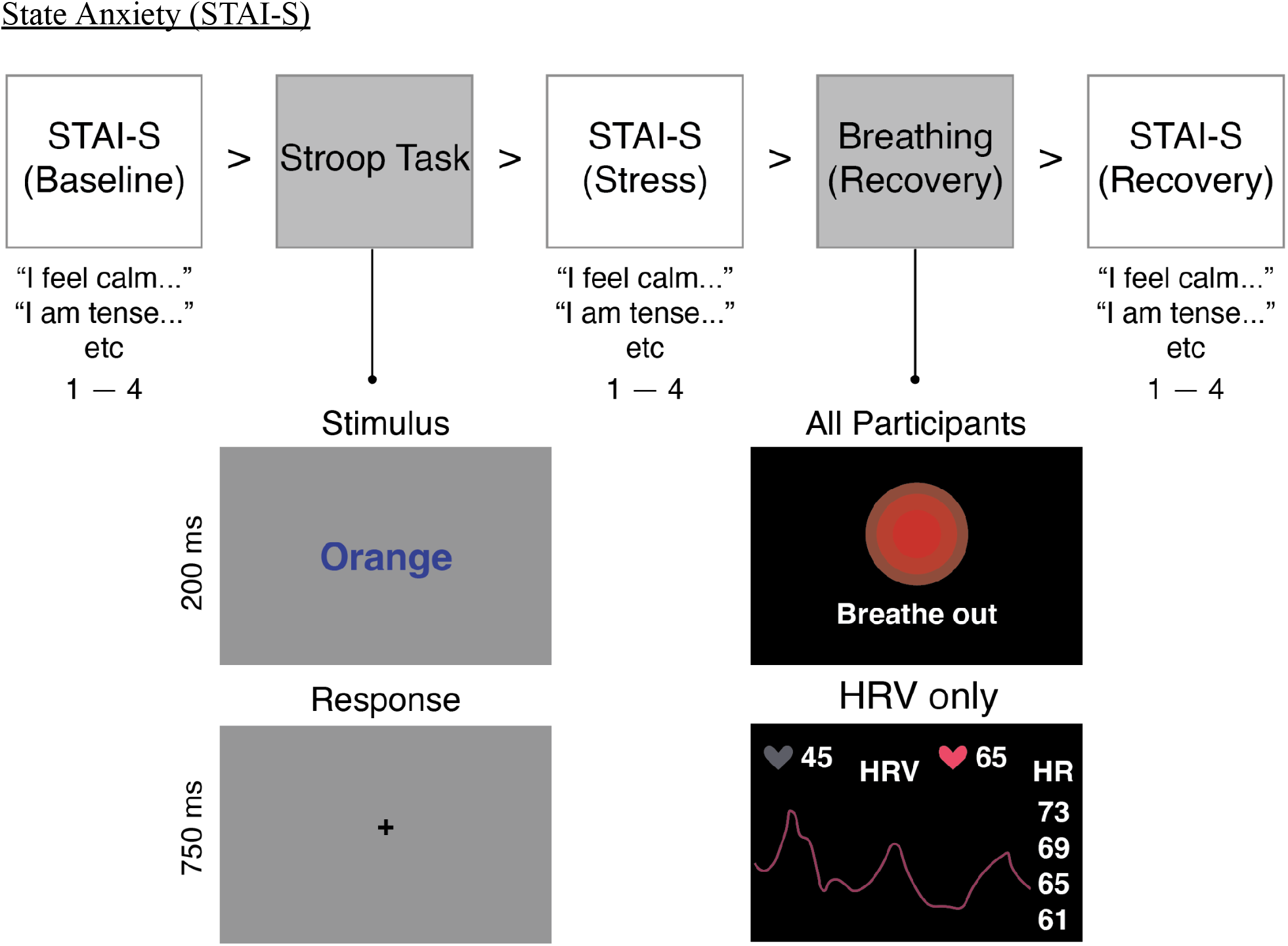
Study design and procedure. Participants completed three STAI-S assessments, a speeded Stroop stress induction, and a 10-minute paced breathing intervention with or without HRV biofeedback. STAI-S was administered before the Stroop task (baseline), immediately afterward (stress), and following the breathing intervention (recovery). Bottom left panel: an example incongruent trial of the Stroop task. Participants were instructed to respond with the color, not identity, of the color-word shown. Bottom right panel: All participants were shown the breathing pacer instructions and circle. Below this, example HRV biofeedback visual display showing real-time HRV trace.

**Figure 2.**
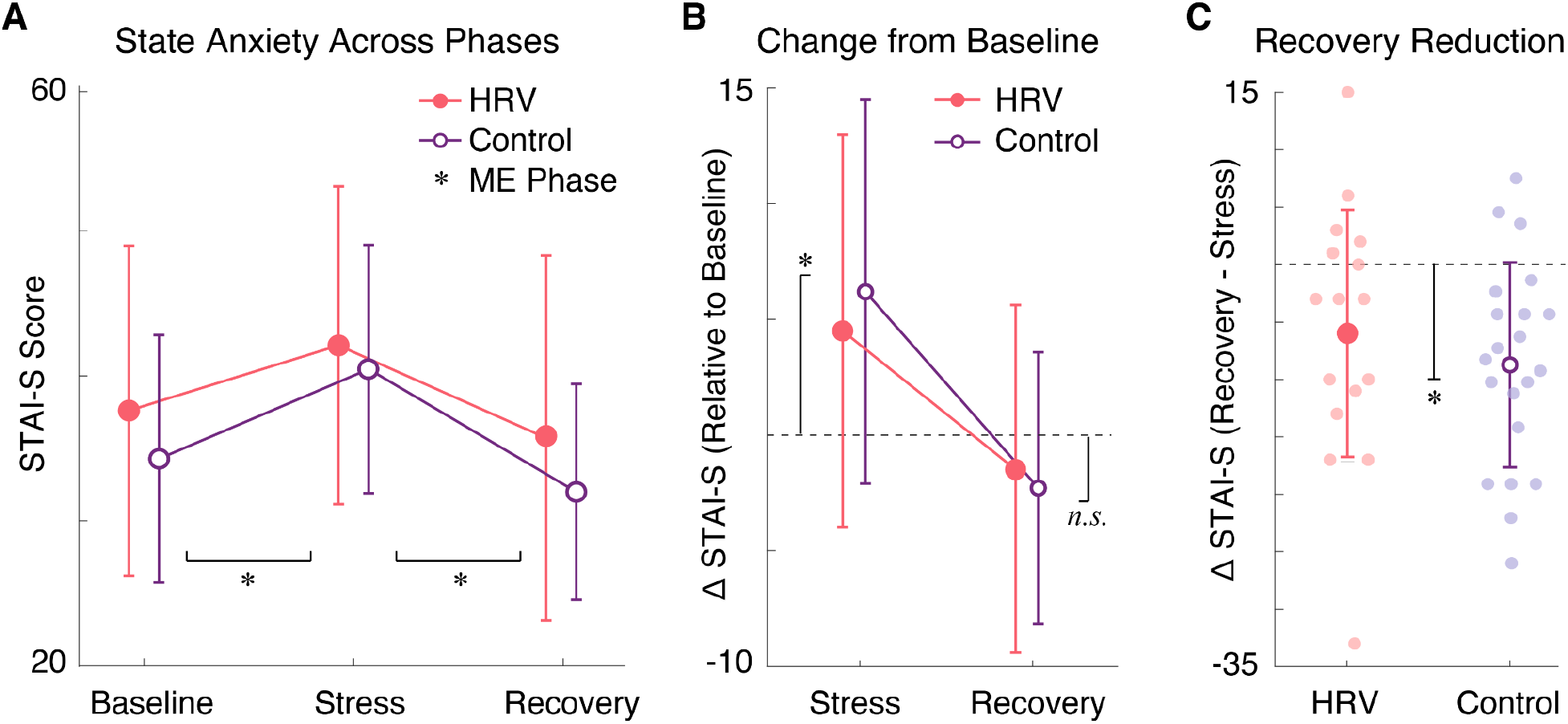
STAI-S outcomes across experimental phases. **(A)** State anxiety scores from baseline through post-Stroop to post-breathing for biofeedback participants (pink) and control participants (purple). Circles show group means. Asterisks denote significant STAI changes between phases at *α* = 0.05 **(B)** Change in STAI-S relative to baseline for biofeedback (pink) and control (purple) participants. Asterisks denote significant STAI-S changes from baseline at *α* = 0.05, dashed line at zero indicates no change from baseline. **(C)** Individual participant STAI-S change scores from post-Stroop to post-breathing, separated by group. As before, means are represented respectively by the solid-filled red circle and outlined purple circle, while the lighter circles in the background represent individual participants. Asterisk denotes significant STAI-S changes from stress to recovery at *α* = 0.05. Dashed line at zero indicates no change from stress. All error bars indicate ± 1 SD.

### EEG Spectral Power: Time-Averaged Effects

We first examined whether EEG power during the breathing intervention differed from baseline across frequency bands, using one-sample t-tests against zero with FDR correction across channels. Across all participants, significant increases from baseline were observed in the alpha and beta bands (Table 1), with alpha increases concentrated over posterior scalp regions (Fig. 3, top row). Changes in the delta and theta bands did not survive FDR correction (Table 1).

**Figure 3.**
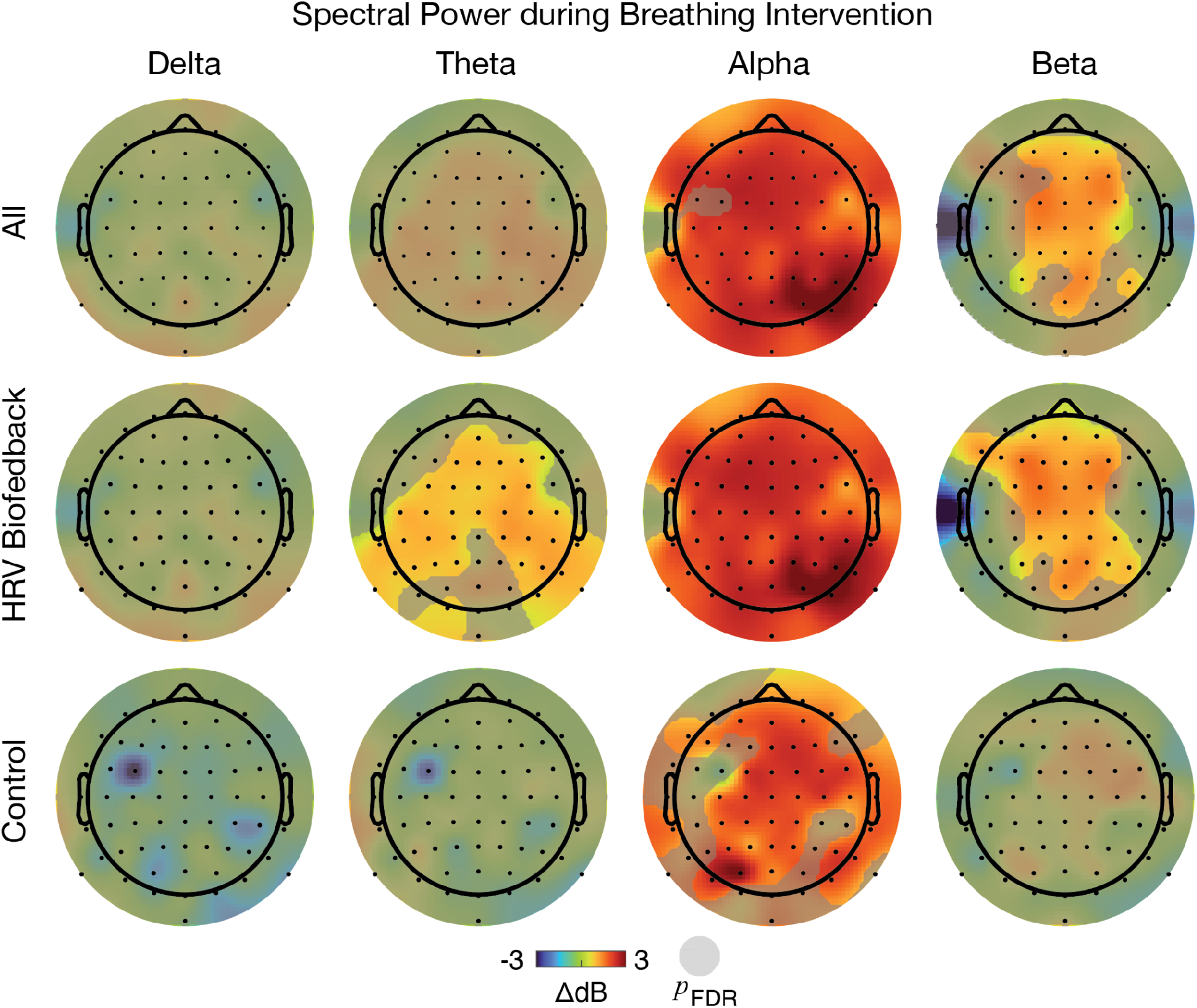
Topographic maps of spectral power change (ΔdB relative to the first 30-second epoch) averaged across the full 10-minute breathing intervention, shown for all participants (top row), HRV biofeedback participants (middle row), and control participants (bottom row). Columns correspond to delta (1–4 Hz), theta (4–8 Hz), alpha (8–13 Hz), and beta (13–30 Hz) bands. Gray overlay indicates electrodes reaching significance in a one-sample t-test against zero after correction for multiple comparisons across channels at *α* = 0.05.

**Table 1.** Topographic statistics in spectral power changes during breathing intervention. Statistics reflect t-tests against zero over all (left column) HRV (middle column) and Control (right column) participants. Asterisks indicate significance after correction for multiple comparisons at *α* = 0.05, *n.s.* denotes that no channels survived significance after correction for multiple comparisons.

| **Band** | **All Participants** | **HRV** | **Control** |
|---|---|---|---|
| Delta | *n.s* *p* > 0.11<br>-1.36 ≤ *t*<sub>36</sub> ≤ 1.64 | *n.s* *p* > 0.008<br>-0.98 ≤ *t*<sub>15</sub> ≤ 3.27 | *n.s* *p* > 0.038<br>-2.30 < *t*<sub>20</sub> < 0.48 |
| Theta | *n.s* *p* > 0.002<br>-1.02 ≤ *t*<sub>36</sub> ≤ 3.45 | \* *p*<sub>FDR</sub> = 0.028<br>2.48 ≤ *t*<sub>15</sub> ≤ 5.32 | *n.s* *p* > 0.08<br>-1.43 < *t*<sub>20</sub> < 1.81 |
| Alpha | \* *p*<sub>FDR</sub> = 0.047<br>2.06 ≤ *t*<sub>36</sub> ≤ 9.18 | \* *p*<sub>FDR</sub> = 0.039<br>2.25 ≤ *t*<sub>15</sub> ≤ 14.10 | \* *p*<sub>FDR</sub> = 0.028<br>2.52 ≤ *t*<sub>20</sub> ≤ 5.39 |
| Beta | \* *p*<sub>FDR</sub> = 0.012<br>2.66 ≤ *t*<sub>36</sub> ≤ 6.25 | \* *p*<sub>FDR</sub> = 0.039<br>-2.25 ≤ *t*<sub>15</sub> ≤ 9.42 | *n.s* *p* > 0.0008<br>-0.99 ≤ *t*<sub>20</sub> < 3.99 |

Post hoc examination of groups revealed pronounced and spatially extensive effects in the biofeedback group. The biofeedback group showed significant alpha increases across a broad distribution of channels compared to controls (Table 1) (Fig. 3, middle and bottom rows). Theta power increased over lateral posterior and central electrodes in the biofeedback group, with no significant modulations in the control group (Table 1). Fronto-central beta power increases similarly survived FDR correction in the biofeedback group but not in controls (Fig. 3). Finally, changes in delta power did not survive correction in any group (Table 1).

### EEG Spectral Power Across Regions of Interest

To characterize how spectral power evolved across the breathing session, we divided the intervention into early (first half) and late (second half) periods (see Methods). Topographic maps computed separately for each period showed that the spatial patterns described above emerged progressively: effects present but weaker in the early period became more spatially robust by the late period (Fig. S2). To directly test for group differences and temporal dynamics, we applied linear mixed-effects (LME) models within posterior and central ROIs (Regions of Interest) using biofeedback condition and time (early vs. late) as fixed factors (see Methods).

We observed greater posterior delta power in biofeedback participants during single session HRV (Fig. 4A) (*F*_1, 70_ = 6.73, *p*_FDR_ = 0.011). Theta power was also elevated in the biofeedback group, in both posterior (*F*_1, 70_ = 6.68) and central (*F*_1, 70_ = 4.52) ROIs (*p_FDR_* = 0.037). We did not observe a main effect of biofeedback in the alpha (*F*_1, 70_ < 1.4, *p* > 0.2) or beta (*F*_1, 70_ < 3.1, *p* > 0.08) bands, in either ROI. This indicates the aforementioned alpha and beta topographic distributions may be too broadly distributed to translate into significant group differences in a more conservative ROI-based analysis.

**Figure 4.**
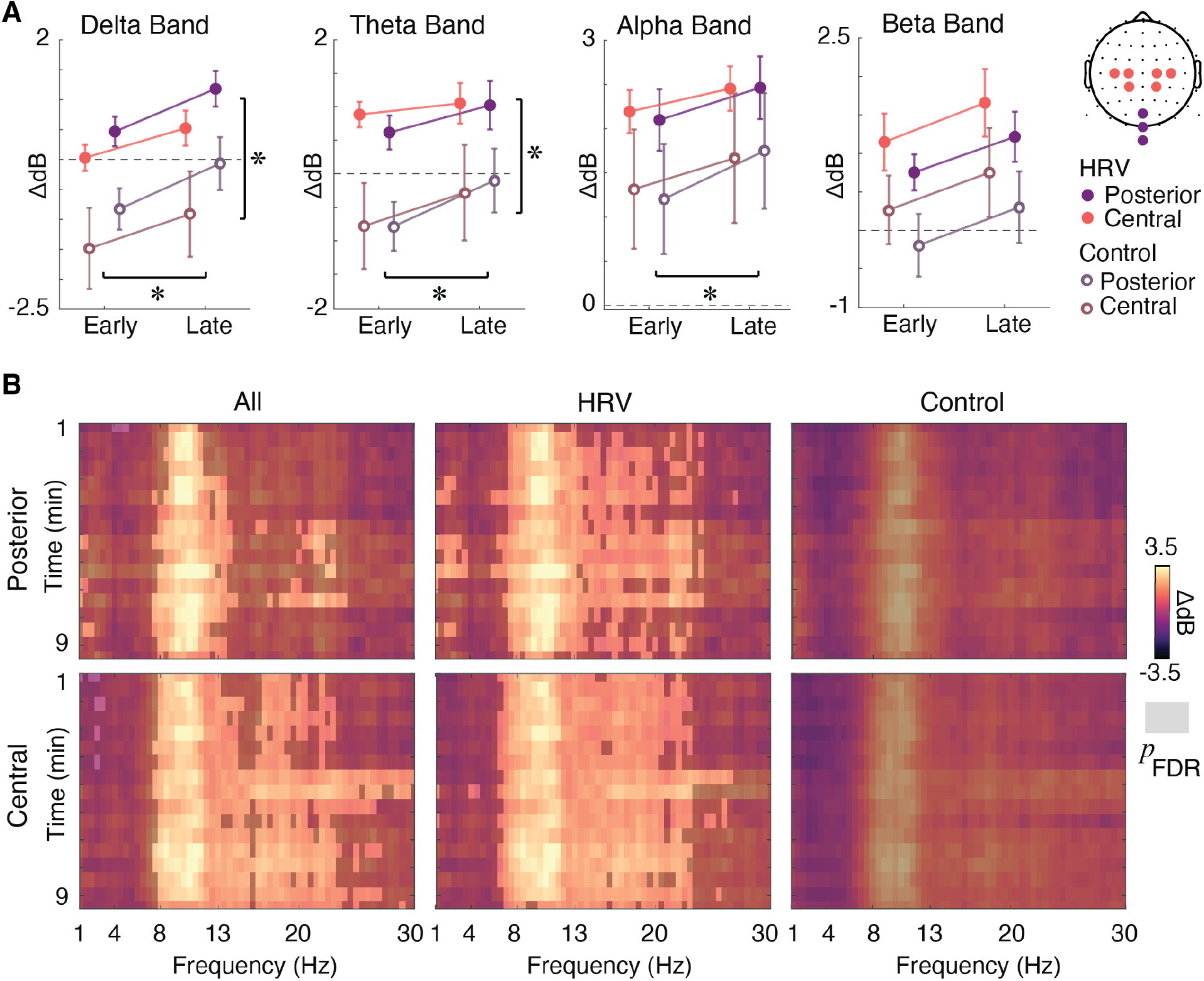
Time-resolved spectral power dynamics during the breathing intervention. **(A)** Band-specific spectral power change (ΔdB) in posterior (purple) and central (pink) ROIs during the early and late phases of the intervention. Filled symbols denote HRV biofeedback participants and open symbols denote controls. Asterisks denote significant main effects of time (horizontal line) and group (vertical line) after FDR correction across ROIs. Error bars indicate ±1 SEM. **(B)** Time-frequency representations of power change relative to baseline for posterior (top row) and central (bottom row) ROIs shown for all participants (left column), HRV biofeedback participants (middle column), and control participants (right column). X-axis shows frequency (Hz); y-axis shows time (minutes from breathing onset). Grey overlay denotes bins significantly different from zero following FDR correction across time-frequency bins at *α* = 0.05.

Spectral power increased from early to late in both ROIs in the delta (posterior: *F*_1, 70_ = 12.52; central: *F*_1, 70_ = 5.56; *p*_FDR_ = 0.021) and theta (posterior: *F*_1, 70_ = 10.71; central: *F*_1, 70_ = 6.60; *p*_FDR_ = 0.012) bands (Fig. 4A). Alpha power increased significantly over time in the posterior ROI (*F*_1, 70_ = 12.52, *p*_FDR_ = 0.018) but not the central ROI (*F*_1, 70_ < 1.7, *p* > 0.2), consistent with the posterior dominance of alpha oscillations. Trends toward a time effect in the beta band did not survive correction (*F*_1, 70_ < 4.1, *p* > 0.04).

No significant group-by-time interactions were observed in any frequency band or ROI (*F*_1, 70_ < 1.2, p > 0.2 in all cases), indicating that the rate of spectral change across the session did not differ between the biofeedback and control conditions. Taken together, HRV biofeedback specifically elevated delta and theta power beyond controls, while increases in delta, theta and alpha power across the session occurred equivalently in both groups (Fig. 4A) — mirroring the pattern of equivalent STAI-S reduction.

### Time-Frequency Analysis

To complement the band-restricted ROI analyses with a model-free characterization of spectral dynamics, we applied one-sample t-tests at each time-by-frequency bin within each ROI. Across all participants, significant modulations from baseline spanned all four frequency bands in both the posterior (*p*_FDR_ = 0.011) and central (*p*_FDR_ = 0.023) ROIs (Fig. 4B) (Table 2). These broad-spectrum effects were replicated in the biofeedback group (Posterior: *p*_FDR_ = 0.021; central: *p*_FDR_ = 0.027) but did not survive FDR correction in the control group (Fig. 4B). This pattern reinforces the conclusion that neural signatures of paced breathing were more robust and widespread with biofeedback, even in the absence of a corresponding advantage in subjective anxiety reduction.

**Table 2.** Time frequency t-statistics against zero corrected across all time frequency combinations within each ROI. Results shown separately over All, HRV and Control (Ctrl) participants, as well as for Delta, Theta, Alpha and Beta frequency bands.

|  | Channel | **Delta** | **Theta** | **Alpha** | **Beta** |
|---|---|---|---|---|---|
| **All** | Posterior<br>*p*<sub>FDR</sub>= 0.011 | -3.17 ≤ *t*<sub>36</sub> ≤ -2.73<br>2.84 ≤ *t*<sub>36</sub> ≤ 3.50 | -3.41 ≤ *t*<sub>36</sub> ≤ -2.71<br>2.70 ≤ *t*<sub>36</sub> ≤ 3.40 | 2.70 ≤ *t*<sub>36</sub> ≤ 5.52 | 2.70 ≤ *t*<sub>36</sub> ≤ 4.52 |
|  | Central<br>*p*<sub>FDR</sub>= 0.023 | -3.43 ≤ *t*<sub>36</sub> ≤ -2.38 | 2.38 ≤ *t*<sub>36</sub> ≤ 3.85 | 2.40 ≤ *t*<sub>36</sub> ≤ 7.19 | 2.37 ≤ *t*<sub>36</sub> ≤ 5.31 |
| **HRV** | Posterior<br>*p*<sub>FDR</sub> = 0.021 | 2.69 ≤ *t*<sub>15</sub> ≤ 4.45 | 2.64 ≤ *t*<sub>15</sub> ≤ 5.47 | 2.61 ≤ *t*<sub>15</sub> ≤ 8.14 | 2.58 ≤ *t*<sub>15</sub> ≤ 5.85 |
|  | Central<br>*p*<sub>FDR</sub> = 0.027 | 2.71 ≤ *t*<sub>15</sub> ≤ 3.28 | 2.46 ≤ *t*<sub>15</sub> ≤ 5.55 | 2.66 ≤ *t*<sub>15</sub> ≤ 11.85 | 2.47 ≤ *t*<sub>15</sub> ≤ 9.38 |
| **Ctrl** | Posterior<br>*p*<sub>FDR</sub><*p* | -3.83 ≤ *t*<sub>20</sub> ≤ 2.78 | -3.69 ≤ *t*<sub>20</sub> ≤ 3.91 | -1.64 ≤ *t*<sub>20</sub> ≤ 3.91 | -2.12 ≤ *t*<sub>20</sub> ≤ 3.06 |
|  | Central<br>*p*<sub>FDR</sub><*p* | -3.26 ≤ *t*<sub>20</sub> ≤ 0.40 | -2.84 ≤ *t*<sub>20</sub> ≤ 1.81 | -0.26 ≤ *t*<sub>20</sub> ≤ 4.25 | -1.07 ≤ *t*<sub>20</sub> ≤ 3.09 |

### Qualitative Reports Reflect Potential Competing Mechanisms

A subset of biofeedback participants verbally reported during or after the session that viewing real-time HRV data felt anxiety-provoking. Of the nine biofeedback participants who completed a brief post-experiment survey on this topic, only three expressed a preference for performing the breathing exercise with the heart rate monitor present. For example, one participant noted that seeing the readout was less relaxing than doing the breathing exercise without it (see Supplementary). We note that these qualitative observations are exploratory and based on a small subsample of the biofeedback group and so should be interpreted with appropriate caution.

## DISCUSSION

The present study examined whether real-time HRV biofeedback enhances the anxiety-reducing effects of paced breathing and whether self-report and neural measures of anxiety relief converge within a single acute session. Two central findings emerge. First, paced breathing reduced subjective state anxiety significantly and equivalently in both groups: both conditions produced full recovery to baseline STAI-S levels after ten minutes, with no HRV biofeedback advantage for. Second, HRV biofeedback produced distinct neural signatures, characterized by greater posterior delta, bilateral theta power, and a broader topographic distribution of alpha and beta power across the full breathing period. Critically, these biofeedback-specific neural effects did not translate into greater subjective benefit. Taken together, these results support the interpretation that paced breathing alone produces acute subjective anxiety relief, while HRV biofeedback induces additional neural changes that, at least within a single-session context, may not benefit the conscious experience of that relief.

### Neural and Self-Report Dissociation

The dissociation between neural and self-report outcomes observed here adds to a growing literature highlighting the imperfect correspondence between subjective anxiety and its physiological indices^36–38^. Fronto-central theta increases in the biofeedback group are consistent with cognitive engagement, working memory and regulatory monitoring^29,39–42^, given their selective presence in participants actively processing the HRV display. This type of cognitive engagement with the feedback signal may limit additional subjective benefit beyond paced breathing alone, is consistent with qualitative reports from biofeedback participants and with prior frameworks proposing an adjustment period before users can productively use real-time physiological information^17^. The present sample had no prior biofeedback experience, and so speaks to the practical question of how biofeedback training should be structured for inexperienced users^29,41,42^.

### Interpretation of Frequency-Band-Specific Effects

The ROI-level analysis found that biofeedback elevated posterior delta and bilateral theta power beyond control levels. Alongside the aforementioned increased cognitive demands of the HRV display, it is also possible that theta increases reflect interoceptive attention specifically^43,44^. Elevated delta activity has been associated with relaxed or internally directed states^45–47^, although its functional significance in acute biofeedback contexts remains unclear. Future work using source localization and connectivity analyses could help distinguish between these accounts.

The significant main effect of time in the posterior alpha band, observed equally in both groups, is consistent with the well-documented role of alpha oscillations in cortical inhibition and relaxed wakefulness^29,41^. The fact that this temporal alpha increase was equivalent in both groups mirrors the equivalent STAI-S recovery, suggesting that slow, rhythmic breathing alone is sufficient to produce progressive posterior alpha increases and subjective anxiety relief.

### Limitations and Future Directions

Several limitations constrain interpretation of the present findings. First, we employed a single-session design; prior literature suggests that biofeedback benefits accumulate with repeated practice^48^ where extended training may attenuate anxiety-inducing effects of the monitoring interface^49^. Second, HRV data were not collected in the control condition, precluding assessment of biofeedback-related modulations in cardiac autonomic regulation. Third, the study used a brief acute stress induction and participants had relatively modest baseline anxiety levels, raising the open question of whether similar dissociations would emerge in individuals with clinical-level or chronic anxiety.

Future work should examine whether repeated biofeedback sessions narrow the gap between neural and subjective outcomes, and whether individual differences in interoceptive awareness, anxiety sensitivity, or prior meditation experience predict the degree of dissociation between these domains. Direct comparisons between HRV biofeedback and other modalities, such as EEG alpha neurofeedback, would also be informative for understanding whether feedback on neural relaxation markers produces greater convergence between neural and self-reported outcomes than autonomic feedback does.

### Conclusions

Ten minutes of paced breathing produced reliable reductions in subjective state anxiety following acute stress induction, regardless of whether HRV biofeedback was provided. Although HRV biofeedback altered EEG spectral dynamics, particularly within delta and theta frequency bands, these neural changes did not translate into additional reductions in self-reported anxiety. These findings suggest that paced breathing alone may be sufficient to produce acute subjective benefit, while the added neural effects of HRV biofeedback may require repeated training or longer-term practice before influencing subjective outcomes. More broadly, the results caution against treating self-report and neural measures as interchangeable indices of anxiety relief in brief, single-session intervention contexts.

## METHODS

### Participants

Forty-eight healthy adults were recruited from the Occidental College community. All were at least 18 years of age and provided written informed consent prior to participation. Participants received monetary compensation. Exclusion criteria included self-reported neurological or psychiatric conditions and current use of psychoactive medication. Participants were randomly assigned to the biofeedback group (N = 24) or the non-biofeedback control group (N = 24). N = 11 participants were excluded from analysis due to issues during the session, including excessive muscle (N = 6) and electrode-noise artifacts (N = 3) and experimenter failure to record EEG data (N = 2). Thus, analyses include N = 37 participants with complete behavioral and EEG data across all three STAI time points (HRV N = 16; Control N = 21). This research was conducted in accordance with the ethical principles of the Declaration of Helsinki, and IRB approval was obtained from Occidental College under protocol FA24-22-DAO.

### Design and Procedure

The experiment comprised three sequential phases: baseline assessment, stress induction, and recovery.

#### Baseline

Following EEG electrode placement, participants completed the STAI-S administered via Qualtrics before completing any tasks.

#### Stress Induction

To elicit acute stress, participants completed a speeded computerized color-word Stroop task (Renaud and Blondin, 1997). A color word (Red, Orange, Yellow, Green, Blue, or Purple) was displayed for 200 ms at the center of a grey screen in one of the six corresponding ink colors, yielding congruent (e.g., the word “Red” in red ink) and incongruent (e.g., the word “Red” in blue ink) trials. The stimulus set was constructed as a fully crossed 6×6 design of word identity and ink color, with each of the 36 conditions repeated six times, yielding 216 trials presented in a randomized order. Each trial began with a variable inter-stimulus interval drawn uniformly from 1–1.5 s. The color word was then displayed for 200 ms, after which participants had a 750 ms response window to press one of six designated keyboard keys (R, O, Y, G, B, P) corresponding to the ink color. Auditory feedback followed each response: a high-pitched tone (800 Hz) indicated a correct response, and a low-pitched tone (100 Hz) accompanied by a red “X” indicated an error or missed deadline. Trials with no response within the deadline were scored as incorrect. Timing was controlled using frame-locked display routines via the MATLAB Psychophysics Toolbox (Brainard, 1997). Following the Stroop task, participants completed the STAI-S a second time.

#### Recovery Paced Breathing Intervention

All participants then completed a 10-minute paced breathing exercise. At the start of this period, the experimenter entered the room to set up the breathing exercise on the iPad. Meanwhile, the computer screen used for the Stroop task displayed the message “Please direct your attention to the breathing exercise” in black text on a plain gray background. Breathing pace was cued visually by a circle that expanded on inhalation and contracted on exhalation, providing a slow, consistent respiratory tempo intended to standardize breathing rate across participants.

The breathing pacer was presented as a single circle, rendered in red, centered on an iPad screen (178.5 mm by 247.6 mm) positioned in front of the participant. The breathing intervention was identical for both the biofeedback and control conditions. At the start of each breath cycle, the circle appeared at its minimum size (1cm diameter), and spiraled outward to four times its starting diameter over a fixed 4-s interval^50^ accompanied by an on-screen “Breathe In” prompt. The circle then spiraled inward back to its minimum size over the subsequent second 4-s interval, accompanied by an on-screen “Breathe Out” prompt. This yielded an 8-s breathing cycle (∼7.5 breaths/min), held constant across the full 10-minute intervention (∼75 cycles). In the biofeedback condition, participants additionally wore a Polar H10 chest-strap heart-rate monitor connected via Bluetooth to the EliteHRV smartphone application^32^, which displayed their real-time HRV with the breathing cue above. Participants in the control condition performed the identical paced breathing exercise using only the visual breathing cue, with no access to physiological data. Following the breathing exercise, all participants completed a final STAI-S administration.

### HRV Display

No participants expressed familiarity with HRV, so all were considered to be novices. In the biofeedback condition, HRV was measured and displayed using the Polar H10 chest-strap monitor (Polar Electro Oy, Kempele, Finland), a research-validated device demonstrating high agreement with reference ECG for time-domain HRV parameters (r > 0.93, ICC > 0.93; Schaffarczyk et al., 2022). The device records RR intervals at 1000 Hz using dry electrodes embedded in a chest strap. The strap was positioned at the level of the xiphoid process and moistened prior to placement to ensure electrode conductivity, consistent with standardized protocols (Flatt et al., 2023). HRV data were streamed via Bluetooth to the EliteHRV smartphone application (Elite HRV Inc., Asheville, NC), which has been validated for accuracy in measuring HRV during slow-paced breathing, with median bias ≤1.3 ms relative to reference ECG (Flatt et al., 2023). The EliteHRV application, displayed on an iPad, provided participants in the biofeedback condition with a real-time graph of their HRV alongside the paced breathing cue^33^.

### Self-Report Measure

State anxiety was assessed using the State subscale of the State-Trait Anxiety Inventory (STAI-S; Spielberger et al., 1983). The STAI-S consists of 20 items rated on a 4-point Likert scale (1 = not at all; 4 = very much so), yielding total scores between 20 and 80. Ten items are positively worded (e.g., “I feel calm”) and ten are negatively worded (e.g., “I feel tense”), with positively worded items reverse-scored before summation. Higher total scores reflect greater state anxiety. A score of 40 or above is commonly used as a clinical threshold for elevated anxiety (Spielberger et al., 1983). The STAI-S was administered at three time points: (1) at baseline, before the Stroop task; (2) immediately after the Stroop task; and (3) immediately after the breathing exercise. All administrations were delivered via Qualtrics.

### EEG Recording and Preprocessing

EEG was recorded using a 64-channel BioSemi ActiveTwo system (BioSemi B.V., Amsterdam, Netherlands), supplemented by eight external electrodes for electrooculographic and mastoid reference recordings. Data were sampled at 1024 Hz.

Offline preprocessing was performed in MATLAB (MathWorks, Natick, MA) using the EEGLAB toolbox (Delorme and Makeig, 2004). Signals were band-pass filtered between 0.1 and 50 Hz using a Butterworth filter and re-referenced to the average of the mastoid electrodes. The PREP pipeline (Bigdely-Shamlo et al., 2015) was applied to remove power line noise at 50 Hz using a multitaper algorithm, identify bad channels, and compute a robust average reference following bad-channel rejection. Independent Component Analysis (ICA) was subsequently applied to remove artifacts associated with eye blinks, eye movements, and other physiological sources; 2.3 ± 1.04 (mean ± SD) components were removed per participant on average. Three individual electrode records were marked as bad post-hoc and excluded from analyses for specific participants (channel 63 for participants 18 and 26; channel 10 for participant 28) based on visual inspection.

### Spectral Analysis

EEG data corresponding to the breathing intervention period were segmented into contiguous 30-second epochs. Spectral power in each epoch was estimated using Welch’s method (MATLAB *pwelch* function) with a 2-second Hanning window, 50% overlap, and an FFT length of max(256, 2^nextpow2(sample rate × 2)) points. This procedure generated power spectra for each channel, and analyses were restricted to the 1–30 Hz range. A total of 19 epochs were computed per participant, encompassing the full breathing interval. Due to recording problems in four participants, the final 30-second epoch was unavailable; it was therefore excluded from all analyses, retaining 19 epochs (1 baseline + 18 post-baseline) across participants.

The first 30-second epoch (beginning at the onset of the breathing task) served as a within-participant baseline. Power in each subsequent epoch was expressed as a decibel (dB) change from baseline using:

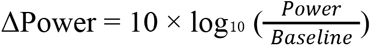

The 18 post-baseline epochs were then averaged into two time bins: an early bin comprising epochs 2–10 (∼minutes 0.5–5 of the breathing exercise) and a late bin comprising epochs 11–19 (∼minutes 5–9.5). Spectral power was analyzed across four canonical frequency bands: delta (1–4 Hz), theta (4–8 Hz), alpha (8–13 Hz), and beta (13–30 Hz).

For topographic analyses, we collapsed spectral power across all 18 post-baseline epochs and applied one-sample t-tests against zero (i.e., against the within-participant baseline) at each of the 64 channels independently for each frequency band. Multiple comparisons across channels were corrected using the false discovery rate (FDR; Benjamini and Hochberg, 1995), applied separately for each band and for all participants, the biofeedback group, and the control group.

For region-of-interest (ROI) analyses, we defined two bilateral electrode clusters based on prior literature and visual inspection of the topographic effects: a posterior ROI comprising electrodes POz, Iz, and Oz, and a central ROI comprising FC1, FC2, FC3, FC4, C1, and C2. Within each ROI, spectral power was averaged across channels and across the relevant frequency bins, then entered into a linear mixed-effects model (LME) using MATLAB’s *fitlme* function:

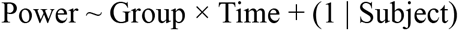

where Group was a fixed between-subjects factor (biofeedback vs. control), Time was a fixed within-subjects factor (early vs. late), and Subject was modeled as a random intercept to account for repeated measures. Statistical inferences were drawn from F-tests obtained via ANOVA on the fitted LME. FDR correction was applied across the two ROIs separately for each frequency band and each effect type (main effect of Group, main effect of Time, and their interaction). Significance threshold was set at *p* < 0.05 after FDR correction.

Finally, an exploratory time-frequency analysis was conducted by applying one-sample t-tests against zero at each time × frequency bin (1–30 Hz, all 18 post-baseline epochs) within each ROI separately. Resulting p-values were FDR-corrected across all time-frequency bins within each ROI and group. This resulted in 18 (time) * 59 (frequency) = 1062 values for correction for each time frequency plot.

### Statistical Analysis

Two primary outcomes were analyzed: STAI-S scores across the three time points and EEG spectral power changes from baseline. For STAI-S, we first tested for group equivalence at baseline using a one-way ANOVA. We then characterized the overall anxiety trajectory across all three time points using a one-way repeated-measures ANOVA, followed by pairwise paired-sample t-tests to examine specific transitions (baseline to post-Stroop; post-Stroop to post-breathing; baseline to post-breathing). Group-by-time effects were tested using a two-way ANOVA with biofeedback condition and measurement time as factors. The magnitude of STAI-S change from post-Stroop to post-breathing was compared between groups using a one-way between-subjects ANOVA. EEG multiple comparison corrections were applied using FDR as described above. Exploratory Pearson correlations were computed to assess the relationship between changes in EEG band power and changes in STAI-S scores across the sample.

## Author Contributions and Funding

CD, IM, KE and SN conceived and designed the study. CD and KE collected the data. SN, KH, CD and IM analyzed the data. SN and CD interpreted the results. SN, CD, and KE drafted the manuscript. SN and CD supervised the project. SN and CD secured funding for the project. This project was supported by a Summer Research Fellowship (2025) awarded to CD; an Academic Student Project grant awarded to CD; and a new faculty startup grant awarded to SN.

