## Supplementary Data for "Single-Session HRV Biofeedback Alters Neural Activity but Not Subjective Anxiety"

Supplementary Results

EEG Analysis

When topographic maps were plotted over time, effects echoed those seen in the results collapsed across time, but with stronger modulations as time increased.

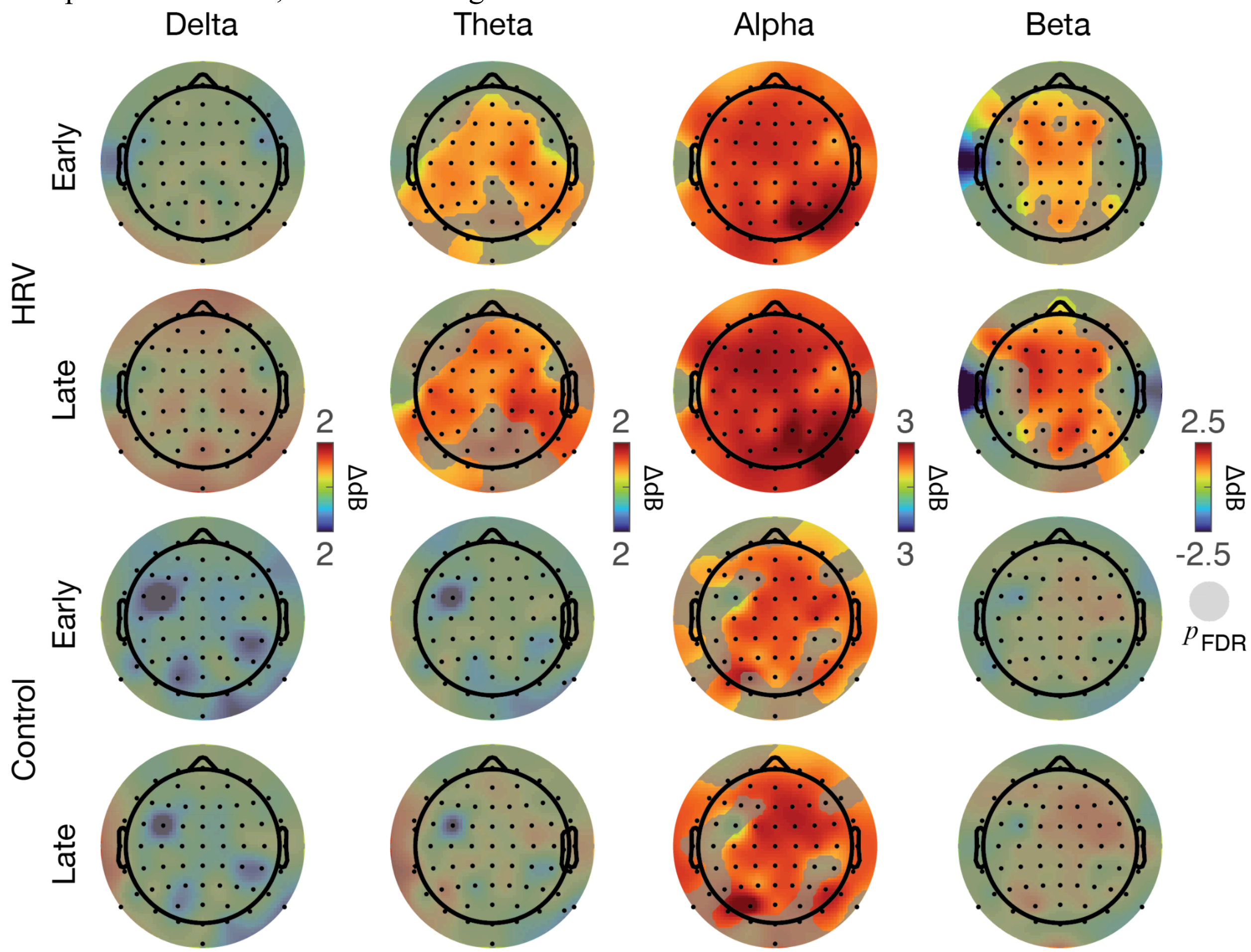


**Supplementary Figure 1.** Topographic maps of spectral power change (ΔdB relative to the first 30-second epoch) averaged across the full 10-minute breathing intervention, shown for HRV participants (top two rows) control participants (bottom two rows). Maps are shown separately for the first (first and third rows) and second (second and fourth rows) halves of the breathing intervention. Columns correspond to delta (1–4 Hz), theta (4–8 Hz), alpha (8–13 Hz), and beta (13–30 Hz) bands. Channel markers indicate electrodes reaching significance in a one-sample t-test against zero after FDR correction at $\alpha = 0.05$.

Qualitative Reports

A post-experimental poll conducted on a subset of biofeedback participants suggest that some may find HRV biofeedback to interfere with stress reduction. As it did not include the entire sample, we use this as supplementary information instead of a main result.

| Respondent | Mental demand | Irritate, discourage or stress | Interfered or helped | Easy to Understand | Affect ability to relax | Prefer to use? |
|---|---|---|---|---|---|---|
| A | 1 | 3 | 4 | 1 | 4 | 3 |
| B | 1 | 2 | 3 | 1 | 3 | 3 |
| C | 1 | 4 | 4 | 1 | 4 | 3 |
| D | 4 | 4 | 4 | 1 | 4 | 3 |
| E | 3 | 1 | 4 | 1 | 4 | 1 |
| F | 3 | 1 | 3 | 2 | 4 | 3 |
| G | 1 | 1 | 1 | 2 | 3 | 1 |
| H | 2 | 2 | 3 | 2 | 3 | 2 |
| I | 3 | 3 | 1 | 1 | 3 | 1 |
| Average | 2.11 | 2.33 | 3.00 | 1.33 | 3.56 | 2.22 |
| STD | 1.17 | 1.22 | 1.22 | 0.50 | 0.53 | 0.97 |

**Supplementary Table 1.** A subset of biofeedback participants completed a post-poll. Abbreviated questions are listed in the header, with average and standard deviation on the last two rows. We show the questions related to the use of the heart rate monitor here; free response as well as full questions and answer options are listed below.

Free response question: Please describe any aspects of the heart rate monitoring that you found particularly helpful, distracting, or challenging:

- Respondent 1: "The heat of the room surged my heart rate so I didn't see it go down at all. Though it wasn't stressful, I was sure that it provided unfavorable results."
- Respondent 2: "I think that wearing the heart monitor and seeing the data made me more aware of my heart rate and how it was changing which kinda in turn freaked me out a little and then cause my heart rate to increase. This made it a little challenging."
- Respondent 9: "Having the heart rate monitor gave something to focus on, but also seeing what my resting heart rate was probably less relaxing then just doing the breathing exercise without it."

Full questions in table:

- How mentally demanding was it to monitor the heart-rate information during the breathing exercise?
    - 1= Not at all, 5 = Demanding
- Did you feel at all discouraged, irritated, or stressed while using the heart-rate monitor during the breathing exercise?
    - 1 = Not at all, 5 = Very
- To what extent did viewing the heart-rate monitor data either interfere with or help your ability to focus on your breathing?
    - 1= Helped, 5 = Interfered
- How easy was it to understand what the heart-rate information was showing you?
    - 1= Confusing, 5 = Easy
- Did viewing your heart-rate data affect your ability to relax?
    - 1= Harder, 5 = Easier
- If you were to complete the breathing exercise again, would you prefer to:
    - 1 = Use, 2 = No preference, 3= Not use
